# Gamma sensory stimulation and effects on the brain

**DOI:** 10.1101/2023.10.30.564197

**Authors:** Martin Kahn, Diane Chan, Danying Wang, Ute Geigenmuller, Cristina Blanco-Duque, Mitchell H. Murdock, Ho-Jun Suk, Brennan Jackson, Vikram Jakkamsetti, Emily Niederst, Emery N. Brown, Edward S. Boyden, Thomas McHugh, Chinnakkaruppan Adaikkan, Annabelle C. Singer, Simon Hanslmayr, Li-Huei Tsai

**Affiliations:** Picower Institute for Learning and Memory, Massachusetts Institute of Technology, Cambridge, Massachusetts, United States of America; Department of Brain and Cognitive Sciences, Massachusetts Institute of Technology, Cambridge, Massachusetts, United States of America; Department of Neurology, Massachusetts General Hospital, Boston, Massachusetts, United States of America, Department of Neurology, Harvard Medical School, Boston, Massachusetts, United States of America; School for Psychology and Neuroscience and Centre for Cognitive Neuroimaging, University of Glasgow, Glasgow G12 8QB, UK; School of Psychology and Centre for Human Brain Health, University of Birmingham, Birmingham B15 2TT, UK; Department of Neurology, UT Southwestern Medical Center, Dallas; Howard Hughes Medical Institute, Cambridge, MA. McGovern Institute, MIT, Cambridge, MA. Departments of Brain and Cognitive Sciences, Media Arts and Sciences, and Biological Engineering, MIT, Cambridge, MA. K. Lisa Yang Center for Bionics, MIT, Cambridge, MA. Koch Institute, MIT, Cambridge, MA; Laboratory for Circuit and Behavioral Physiology, RIKEN Center for Brain Science, Wakoshi, Saitama 351-0198, Japan; Center for Brain Research, Indian Institute of Science, Bangalore, India; Coulter Dept. of Biomedical Engineering. Georgia Tech & Emory University, Atlanta, GA

## Abstract

Findings by the Tsai lab and others ^1–8^ demonstrate that 40 Hz frequency sensory stimulation induces electrophysiological responses and attenuates pathology in mouse models of Alzheimer’s disease (AD). A recent study in *Nature Neurosciene* ^9^ concluded that the stimulation does not affect endogenous gamma oscillations or amyloid burden. We welcome research investigating 40 Hz sensory stimulation, and the article by Soula et al enhances our understanding of the brain’s electrophysiological response to 40Hz. However, we respectfully suggest that the data in Soula et al are consistent with a neuronal response to 40 Hz, which we further support with new data in humans. Moreover we contend the non-significant effects on amyloid are due to technical limitations of the study.

---

In particular we note: 1) A key conclusion from Soula et al. is that the 40 Hz brain response may not directly involve increased <u>endogenous</u> gamma rhythms, but this claim may reflect a semantic disagreement over the word “entrainment” as several lines of evidence, including from Soula et al., reveal a 40 Hz neural response following sensory stimulation. 2) Human data show that 37-40 Hz sensory stimulation causes a neural LFP response in cortical and subcortical structures. 3) Soula et al. do not observe amyloid reduction following 40 Hz sensory stimulation likely due to technical differences in measurement approaches, animal models, and statistical power of the experiments.

## 40 Hz electrophysiological response throughout the brain

Soula et al. and others^10^ question whether the brain’s response to 40 Hz stimulation constitutes “entrainment.” This contention may stem in part from diverging semantic definitions of “entrainment.” In our studies, “entrainment” describes how externally applied 40 Hz stimulation causes a 40 Hz response in the brain. However, entrainment can also describe how an external rhythmic stimulus synchronizes *internal native* oscillations^11^. By this definition, 40 Hz sensory stimulation only “entrains” the brain if it affects endogenous gamma rhythms. We acknowledge that this is a relevant distinction for understanding the neural mechanisms of sensory entrainment. We hope further research will be inspired by our discussion of Soula et al. in the light of previous work and new human data presented herein.

A long history of neuroscience--including Nobel laureate Edgar Adrian’s work in the early 1900s- -reveals that external sensory stimulation induces a rhythmic neural response (e.g., see^12^). Our work has described neuronal spiking and local field potentials (LFP) in response to sensory rhythmic stimulation. These measurements reflect different aspects of activity, and it is unknown whether the effects on AD pathology from 40 Hz rely on synaptic signaling or local neuronal spiking. Our first study in AD mouse models recorded LFP and spiking responses to unimodal 40 Hz visual flicker in the primary visual cortex (V1) and hippocampus of 5xFAD mice^2^. Neural responses to 40 Hz light stimulation in other regions in C57BL/6 mice^1^ were examined and found to increase 40 Hz power in the LFP of V1, hippocampus, somatosensory cortex, and frontal cortex. In the single unit spiking response of hippocampal CA1, we found an increase in phase-locking compared to a control condition where light was occluded^1^.

Soula et al. recorded single units and LFPs from V1, CA1 and the entorhinal cortex (EC) in response to 40-42 Hz^9^ visual stimulation. There are several considerations that should be applied to the data:

1. Soula et al. observe a narrow 40 Hz LFP peak. This is consistent with other work in humans and mice and suggests that the circuit mechanisms of 40 Hz stimulation may have limited overlap with endogenous gamma. We look forward to more research on this question.
2. The authors state that *“Our findings with 40-Hz stimulation can be contrasted with the effect of 4-Hz, high-contrast stimuli [*…*]”* These data are from a public dataset that uses different mice, different types of electrodes, different stimuli and a different stimulation apparatus. Thus it is difficult to compare these datasets.
3. The lack of hippocampal LFP response contrasts with our data and others in humans and mice (Figure 1) and should encourage further research. Of note, the LFP can reflect synaptic inputs, which are a prerequisite for the modulation of CA1 spike timing, as is observed in Soula et al.
4. When discussing neuronal phase locking to 40Hz stimulation, the authors report phase-locking in “a substantial fraction (28%)” of V1 neurons ” but only a “small fraction” of neurons in CA1 (6.7%). We would like to highlight two considerations: First, supplementary figure 4 in Soula et al shows that 15% of CA1 neurons classified as interneurons show significant modulation by 40Hz stimulation. In our view this is an interesting finding worth highlighting, especially given that interneurons are key signaling partners for non-neuronal cells (e.g. vasculature). Second, it is hard to say what fraction of modulated cells is biologically meaningful. Indeed, Soula et al only quantified modulation at the spiking level, which means a substantially larger fraction of cells receive synaptic inputs by 40 Hz stimulation.
5. The study by Soula did not consider the behavioral state of the mouse, described as “*awake but mainly immobile*”. Adaikkan et al. only analyzed periods where mice were moving. As shown in **Figure 1 C-D**, the brain/cognitive state in humans modulates the 40 Hz response, and this may be also true in mouse and explain some of the differences in electrophysiological data between Soula and Addaikan.

**Figure 1.**
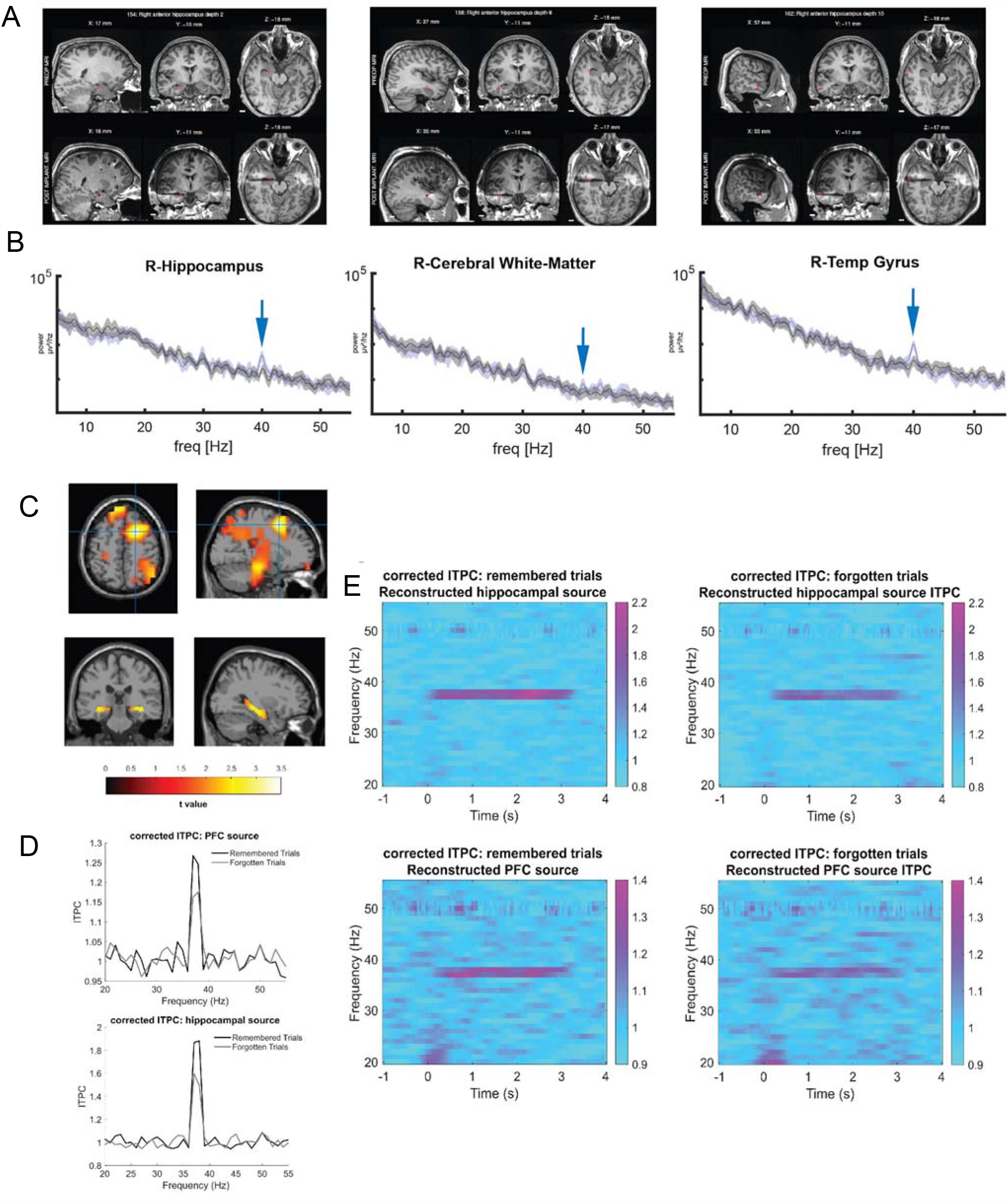
Human brain response to 40 Hz light stimulation. **A)** MRI images showing positions of recording electrodes in the right hippocampus, white matter, and right temporal gyrus. **B)** Power spectral density after Gram-Schmidt correction during a 40 Hz light stimulation (blue) or control stimulation (black) in electrodes shown in A. Solid lines are mean for one subject and shading represents bootstrapped 95% confidence intervals. N=4 recordings from 1 subject. **C-E**: Differences in corrected intertrial phase coherence (ITPC) during encoding between subsequently remembered trials and subsequently forgotten trials. **C)** The ITPC was averaged between 0.5 and 2.5 s and between 37 and 38 Hz. The paired-sample t statistics between the averaged ITPC in the remembered trials and forgotten trials across participants (N = 48) was interpolated to the template MRI. Three significant positive clusters were found when a non-parametric randomization test was restricted to the hippocampus (*p* < 0.05). Two significant positive clusters were found when restricting the ROIs to the left and right superior frontal gyri, the left and right middle frontal gyri and the left and right inferior frontal gyri. **Top:** The strongest ITPC difference was localized, MNI coordinates: 20, 11, and 50. **Bottom:** The t values were masked by the hippocampal ROIs (AAL atlas). **D) Top:** The grand average ITPC (N = 48) was computed for the left and right superior frontal gyri, the left and right middle frontal gyri and the left and right inferior frontal gyri and averaged across 0.5 and 2.5 s and across these channels for each condition, plotted as a function of frequency. **Bottom:** Same as **Top** but the ITPC was computed and averaged across the channels in the left and right hippocampi. The cluster-based randomization test was done to the averaged data (37 - 38 Hz). Significant differences were found when restricting the ROIs to the hippocampus and the PFC. Then the power spectrum was averaged across the virtual channels in each ROI to show the peak over 37.5 Hz.. **E) Top:** Time frequency representations of the grand average ITPC (N = 48) at the hippocampus for the remembered trials (left) and for the forgotten trials (right). **Bottom:** Same as Top but the time frequency representations are the mean across the PFC channels.

## Humans show neural response to gamma sensory stimulation

Mice and humans rely on different sensory modalities ^13^ and have differences in sensory integration ^14^, so it is important to consider the response of sensory flicker in humans. Visual or audio-only stimulation in humans shows a detectable effect size of rhythmicity ^11,15,16^. Moreover, several groups have now stimulated humans with gamma-range visual and/or audio stimulation and recorded significant brain responses, even in deeper brain regions ^17–22^. These results suggest the human brain induces a gamma-range neural response to gamma-range sensory input. Below, we describe experimental evidence highlighting a deep brain region response to gamma sensory stimulation, further highlighting the effect of noninvasive sensory stimulation on human neural activity and behavior.

In tests performed at University of Iowa and analyzed at MIT ^23^, volunteers with stereotactically-implanted depth electrodes were exposed to 40 Hz light stimulation conditions for one minute, followed by one-minute long baseline recordings when the device was on but the stimulation was obscured. Intracranial EEG recordings from hippocampus and cortex (**Figure 1A, B**) of one patient show power at 40 Hz in the spectral density after Gram-Schmidt correction^24^, when shown the 40 Hz light stimulation alone, versus a light-obscured control.

A second group at University of Glasgow independently performed audio/visual stimulation at 37.5 Hz with healthy human volunteers. Forty-eight participants viewed videos and listened to sounds (both flickering/fluttering at 37.5 Hz) and were asked to remember the associations between them. Later, 24 participants were cued with one of the sounds and recalled the paired video. The other 24 participants were cued with a video and asked to recall the paired sound ^22^. The results (**Figure 1C-E**) not only show phase-locked responses in the hippocampus and prefrontal cortex (PFC) to the stimulation, but they also show that this response is modulated by memory success (i.e., stronger responses to remembered vs. forgotten trials)^22^. This suggests the responsiveness of higher-order downstream regions to sensory rhythmic stimulation is modulated by the cognitive state of the brain (e.g. attention).

## Cellular effects of stimulation evoked responses

Previous findings, by our lab and others, have shown effects of 40 Hz stimulation on a wide array of phenotypes in neurodegenerative disease, including protein clearance, synaptic potentiation, autophagy, glial activation, inflammatory changes, and vascular responses ^1– 3,7,8,25,26^. These effects may be indirect from the neural response to gamma stimulation, but it is unclear how neural response to sensory stimulation is coupled to biological pathways related to pathology. Disentangling these questions is an active area of investigation.

Soula et al. investigated attenuation of amyloid burden following noninvasive 40 Hz light stimulation using unconventional experimental methods. The authors pooled results from multiple AD mouse models, sexes, and ages, making the comparison of amyloid load virtually impossible. For example, Soula et al. observe comparable amyloid levels between 4- and 12 - month old mice but it is widely reported that amyloid burden increases over this range in these models^27–29^. Additionally, amyloid was measured by ELISA analysis on PFA-fixed tissue, yet no paper we know of in AD literature has attempted to quantify amyloid using ELISAs on fixed tissue, as the assay is extremely sensitive to protein degradation and conformation. The authors also lacked a positive control to show their experimental approaches detect changes in amyloid beta.

We encourage efforts to investigate the mechanisms of noninvasive sensory stimulation as a potential therapeutic intervention, and the use of rigorous methods, thorough controls, and appropriate analytical approaches.

## Supplemental Methods

### Acquisition of human data

Full details on the acquisition of human data can be found in ^23^ and ^22^. New analyses were run on our previously published data.

The part of the Phase 1 study (NCT04042922, ClinicalTrials.gov) conducted at the University of Iowa was approved by the University of Iowa Institutional Review Board. These studies were carried out in accordance with the Code of Ethics of the World Medical Association. Written consent was obtained by study personnel from all participants. Full inclusion information can be found at https://doi.org/10.1371/journal.pone.0278412.s009. The study record is https://clinicaltrials.gov/study/NCT03702127

Intracranial EEG was recorded via stereotactically-implanted depth electrodes and grid arrays were implanted in medically intractable epilepsy patients for other medical purposes as described in detail in^23^. With the approval of the Committee on the Use of Humans as Experimental Participants at the University of Iowa and in collaboration with Aaron Boes’ group, these studies were done in accordance with the Code of Ethics of the World Medical Association. Written consent was obtained by study personnel from all participants. These studies were done to evaluate for induced entrainment as a result of sensory stimulation. Participants were exposed to three 40Hz stimulation conditions (light alone, sound alone, synchronized light and sound) for 1 minute each interspersed with 1 minute long baseline recordings when the device was on but the light was obscured.

Intracranial EEG was recorded during the entire stimulation experiment using electrode arrays (Ad-Tech Medical Instrument) that included stereotactically-implanted depth electrodes (4-8 macro contacts per electrode, spaced 5-10 mm apart) and grid arrays (containing platinum-iridium disc contacts, with 2.3 mm exposed diameter, 5-10 mm inter-contact distance, embedded in a silicon membrane) placed on the cortical surface. A subgaleal electrode was used as a reference. Data acquisition was controlled by a TDT RZ2 real-time processor (Tucker-Davis Technologies). Collected data were amplified, filtered (0.7-800 Hz bandpass, 12 dB/octave roll-off), digitized at a rate of up to 20 kHz, and stored for subsequent offline analysis.

For scalp EEG data acquired at University of Glasgow, participants in both the visual and auditory cue groups provided informed consent and were given task instructions, before practicing the procedure with four example trials.

### Analysis of 40Hz human stimulation data

To obtain the PSD, the signal was high-pass filtered at 2 Hz and then re-referenced using the Gram-Schmidt process^24^ to subtract the average signal of all other electrodes whilst avoiding the introduction of a spurious 40 Hz signal measured in other electrodes (see^24,30^). The signal was then subdivided into 2s long segments and spectral power in each segment was calculated using Welch’s method. To obtain 95% confidence intervals, the average across all segments was bootstrapped 10000 times.

### Analysis of 37.5 Hz stimulation data

The scalp EEG data during encoding phase of all channels were transformed to scalp current density (SCD) using the finite-difference method (Huiskamp, 1991; Oostendorp & Oosterom, 1996ref.). The leadfield was also SCD transformed by applying the transformation matrix that was used for transforming the potential data (Murzin et al., 2013ref.). Source analysis was done using a linearly constrained minimum variance (LCMV) beamforming method (Van Veen et al., 1997ref.), resulting in 2020 virtual electrodes for each participant. Time-frequency analysis was applied for each trial at each virtual electrode using multitaper time-frequency transformation based on Slepian sequences as tapers. The complex Fourier-spectra was computed using 1s sliding time window for frequencies of interest between 20 and 55 Hz (1 Hz width of smoothing frequency) and time of interest between -1 and 4 s. The complex Fourier-spectra was then normalised by its magnitude. The ITPC was computed by taking the magnitude of the mean of the normalised complex Fourier-spectra across subsequently remembered trials and across subsequently forgotten trials, respectively. ITPC was corrected by a Rayleigh Z correction with n x ITPC^2^ to count the contribution of trial numbers (n = number of trials in each condition) (Michelmann et al., 2019; Samaha et al., 2015ref.), although the trial numbers did not differ statistically significant between remembered and forgotten conditions.

A two-tailed paired-samples permutation test with the Monte Carlo method of 1000 randomisations was conducted between ITPC in the remembered and in the forgotten trials by masking the ROIs including the left and right hippocampi. The same analysis was applied to the ROIs including the left and right superior frontal gyri, the left and right middle frontal gyri and the left and right inferior frontal gyri.

### MRI

MRI data were acquired using a 3-Tesla Siemens Tim Trio scanner (Siemens, Erlangen, Germany) paired with a 12-channel phased-array whole-head coil. Head motion was restrained with foam pillows. Sequences included 3D T1-weighted magnetization prepared rapid acquisition gradient echo (MP-RAGE) anatomical images. To allow for T1-equilibration effects, 4 dummy volumes were discarded prior to acquisition. Online prospective acquisition correction (PACE) was applied to the EPI sequence. Structural MRI data was analyzed using FSL (https://fsl.fmrib.ox.ac.uk/fsl) and FreeSurfer (http://surfer.nmr.mgh.harvard.edu).

## Data Availability Statement

Human intracranial EEG data from the MIT/University of Iowa can be found, deidentified, at https://dataverse.harvard.edu/dataset.xhtml?persistentId=doi:10.7910/DVN/56XZ3F

The scalp EEG data from University of Glasgow can be found here:

- The datasets are available from: https://osf.io/fpyqk/.
- All original code has been deposited at https://osf.io/fpyqk/

—Any additional information required to reanalyze the data reported in this paper is available from the lead contact upon request

## Author contributions

M.K.- conceptualization, iEEG analysis, writing. D.C.- iEEG data acquisition, writing. D.W.- conceptualization, EEG acquisition and analysis. U.G.- iEEG analysis. CBD- writing. MHM- writing. HJS- iEEG data acquisition. BJ- iEEG data acquisition. VK- iEEG analysis. EN- conceptualization, writing. ENB- conceptualization. ESB- conceptualization.TM- conceptualization, writing. CA- conceptualization, writing, ACS- conceptualization, writing. SH- cosupervised, funding, writing. LHT- cosupervised, funding, writing.

